# Effect of membrane composition on interaction and permeation of a mycobacterial siderophore

**DOI:** 10.1101/2025.03.27.645699

**Authors:** Fatemeh Sharifian, Mina Ebrahimi, Ahmad Reza Mehdipour

## Abstract

Iron acquisition is a crucial determinant of Mycobacterium tuberculosis (Mtb) survival and pathogenesis, as the bacterium must scavenge iron from the host environment during infection. To achieve this, Mtb synthesizes two structurally distinct siderophores, mycobactin (MBT) and carboxymycobactin (cMBT), efficiently chelate iron and facilitate its uptake. While MBT remains embedded in the membrane due to its hydrophobicity, cMBT is secreted to scavenge extracellular iron. Despite their biological importance, the dynamics of siderophore–membrane interactions remain poorly understood. In this study, we employ coarse-grained molecular dynamics simulations to investigate the spontaneous flip-flop of iron-bound MBT (Fe-MBT) across the mycobacterial membrane. Our results reveal that Fe-MBT flip-flop occurs on a significantly slower timescale compared to generic bacterial lipids, with rates nearly an order of magnitude lower. The lipid composition of the membrane plays a critical role in modulating this process, with mycobacterial membranes exhibiting a higher degree of lipid order and reduced Fe-MBT mobility compared to generic bacterial membranes. Furthermore, we show that flip-flop transitions are relatively fast, occurring within 100 ns and, in some cases, on the order of a few nanoseconds. Our findings contribute to a mechanistic understanding of siderophore dynamics in Mtb membranes and provide a foundation for future studies on iron acquisition and potential drug targeting strategies.

## 2 Introduction

Iron deprivation represents a critical environmental stressor that *Mycobacterium tuberculosis* (Mtb) encounters during infection, as free iron is unavailable in the human host [1]. To overcome this challenge, Mtb produces two species of siderophores—mycobactin (MBT) and carboxymycobactin (cMBT)—small iron-binding molecules that scavenge iron from the host [2]. These siderophores are exported across the inner membrane via the membrane proteins MmpS4, MmpS5, MmpL4, and MmpL5 [3, 4, 5]. Under low iron conditions, MBT and cMBT efficiently chelate ferric iron, primarily by extracting it from transferrin, before transporting it into the bacterium [6].

Structurally, Mtb’s siderophores are classified as hydroxamate and mixed-type siderophores [5]. MBT and cMBT share similar structures, differing mainly in their lipid residues [7]. While MBT is a lipophilic, intracellular siderophore embedded in the membrane, cMBT is hydrophilic and functions extracellularly. Given their crucial role in iron acquisition, these siderophores are of significant pharmacological interest as potential drug targets for tuberculosis treatment [5]. In fact, siderophore analogs have already shown promise as antibacterial agents [5].

Numerous studies have sought to characterize the structure and function of Mtb’s siderophores [8]. The structures of MBT and its derivatives have been resolved in both their free and metal-bound states [9, 10]. MBT biosynthesis is nonribosomal peptide synthetase-dependent [11]. Both MBT and cMBT exhibit a high affinity for *Fe*^3+^ with an estimated binding constant (*K*_*a*_) of 10^52^ *M*^−1^ [5]. These siderophores share a common chemical core, which consists of a 2-hydroxyphenyloxazoline moiety linked by an amide bond to an acylated N-hydroxylysine residue esterified at the α-carboxyl group with a β-hydroxy acid. The, β-hydroxy acid is attached via an amide bond to a seven-membered lactam ring, and *Fe*^3+^ is coordinated by the hydroxamic acid groups of the N-hydroxylysines, the phenolate oxygen, and the nitrogen atom of the oxazoline moiety (Figure 1).

**Figure 1:**
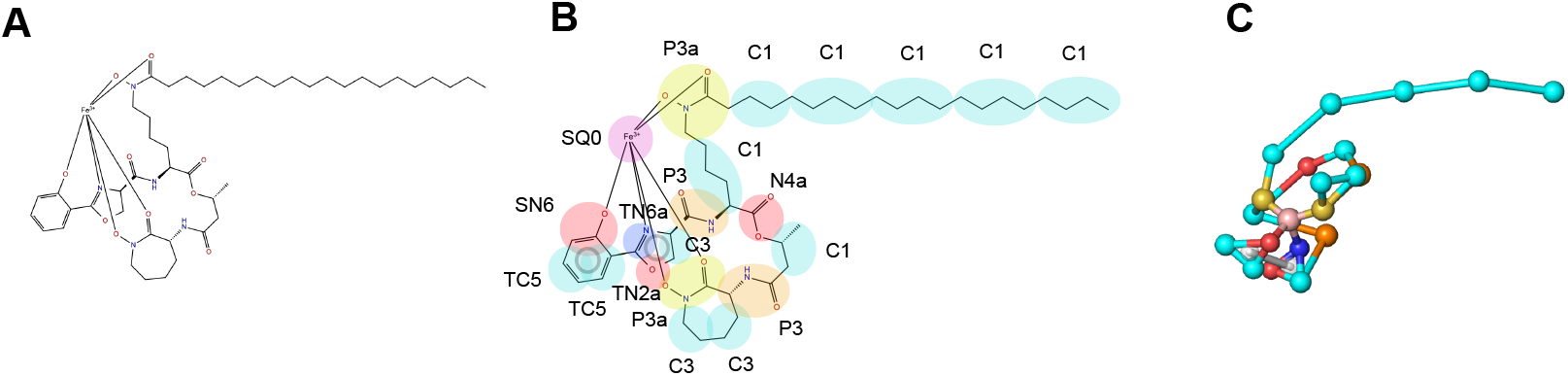
Structure of Fe-mycobactin T (Fe-MBT). **A**) Atomistic representation of Fe-MBT structure. **B**) CG mapping of Fe-MBT. **C**) Martini 3 CG model of Fe-MBT.

Despite their structural similarities, MBT and cMBT differ in their hydrophobicity and cellular localization [5]. With its long aliphatic tail, MBT is hydrophobic and embedded within the membrane. In contrast, cMBT has a shorter tail with a carboxylate group, making it hydrophilic. MBT chelates ferric iron within the plasma membrane on the periplasmic side, where it is likely transported across the membrane by the heteromeric ABC transporter IrtAB, possibly via a sliding scramblase-like mechanism [12, 13]. However, the precise details of this transport process remain unclear.

The complexity of Mtb’s cell envelope poses significant challenges to studying these processes. The Mtb cell envelope consists of four layers: the mycomembrane, which contains mycolic acid esters and other complex lipids; the arabinogalactan-peptidoglycan layer; the periplasmic space; and the plasma membrane [14]. The plasma membrane plays a critical role in regulating the uptake of small molecules and ions. It is composed of various (glyco)lipids, including cardiolipin (CL), phosphatidylethanolamine (PE), phosphatidylmyoinositol (PI), and phosphatidylmyoinositol mannosides (PIMs) [15]. PIM lipids, which are modified forms of PI, contribute to the membrane’s structural diversity and asymmetry. These lipid complexities make it difficult to probe the dynamics of hydrophobic molecules within the membrane.

To address these challenges, coarse-grained molecular dynamics (MD) simulations have become a valuable tool for studying the interactions between proteins, lipids, and small molecules [16]. Coarse-graining simplifies the system by grouping atoms into larger particles, allowing simulations over longer timescales. These are crucial for observing phenomena that occur too slowly to be captured in atomistic simulations [17]. This is especially valuable for studying processes such as the spontaneous flipping of molecules like siderophores between leaflets of the bilayer, which are notoriously slow and difficult to capture in atomistic simulations [18]. Additionally, coarse-grained simulations make it possible to explore the dynamics of complex biological membranes, such as the mycobacterial plasma membrane, which are often too computationally demanding for atomistic simulations due to slow convergence and the large size of the systems involved. Using coarse-grained simulations, we can probe larger-scale behaviors and interactions within these complex environments that would be computationally expensive or even infeasible using an atomistic approach. This allows us to gain insights into membrane dynamics, lipid bilayer asymmetry, and the influence of membrane properties on siderophore transport processes, all of which are essential for understanding the iron uptake mechanisms in *Mycobacterium tuberculosis*. However, specific force-field parameters are required for all system components, including proteins, lipids, and small molecules, to simulate these interactions accurately. While coarse-grained force fields like Martini have been widely used and expanded, they typically do not cover metal-bound complexes, requiring the development of custom parameters for such systems.

In this study, we have developed MARTINI-based coarse-grained force field parameters for iron-bound MBT (Fe-MBT). With these parameters, we investigated the interaction of the Mtb siderophore with the biological membrane at a coarse-grained level. This enabled us to explore longer timescales and gain insights into processes that are otherwise difficult to observe at the atomistic level.

## 3 Methods

### 3.1 Development of MARTINI-based coarse-grained force-field parameters

For the parametrization of the ligands in the MARTINI force field, we adhered to the guidelines outlined for the MARTINI 3 models described by Alessandri et al. [19]. These guidelines include several key steps to ensure that the coarse-grained (CG) representation of the siderophores is accurate and robust. First, we generated a suitable mapping of the ligands by simplifying their atomistic structure into fewer interaction sites, which is a crucial step for coarse-graining. This mapping is designed to maintain the essential features of the molecular structure while reducing computational complexity (Figure 1).

Next, we assigned bead types to each ligand based on their chemical nature and the properties of the atoms they represent. The MARTINI force field uses predefined bead types for different chemical groups, such as hydrophobic, hydrophilic, and charged residues. For the siderophores, this step involved identifying the appropriate bead types for each molecule component, such as the hydroxamate groups, hydrophobic tails, and the coordination sites for the iron atom. The bead type assignments for Fe-MBT are shown in Figure 1, where the CG beads are color-coded according to their size, which reflects their interaction potential in the coarse-grained model.

The third step used a bottom-up approach to parameterize the bonded terms (such as bond lengths, angles, and dihedrals). This step was based on reference atomistic simulations of the ligand placed in a POPE membrane, where we utilized the CHARMM36 force field to generate the necessary data. Atomistic simulations provided high-fidelity data on the behavior of the ligands at the molecular level, which were then mapped to the coarsegrained representation. This approach ensures the CG model retains the original atomistic system’s important structural and energetic properties (Figure S1 & S2).

For model validation, we first checked the solvent-accessible surface area (SASA) of the ligand in the coarse-grained model and compared it with atomistic reference data. This step ensures that the coarse-grained model accurately reflects the solvent-exposed surface of the ligand, which is essential for predicting how the siderophores will interact with the biological membrane (Figure S3) [20].

#### 3.1.1 Coarse-grained molecular dynamics simulations

Using the modified scripts from the CHARMM-GUI web server, the ligands were placed in heterogenous bilayers comprising different lipid compositions [21]. The list of simulations is tabulated in Table 1. Sodium and chloride ions were added to make the ion concentration 150 mM. The Martini 3 force field was used for lipids, ions, and water molecules in coarsegrained simulations [22].

**Table 1:**
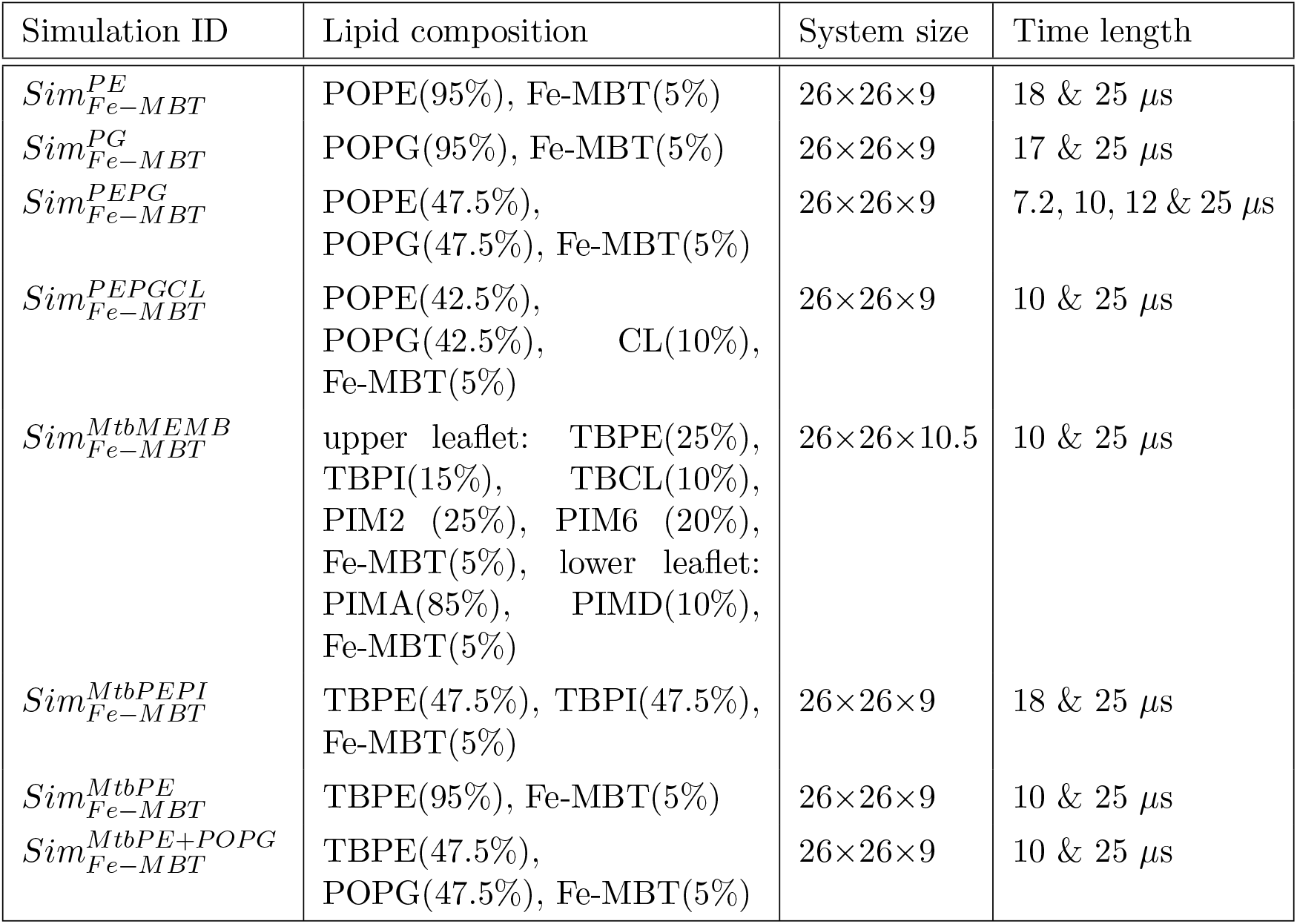
List of different simulations and their lipid composition.

We performed all simulations using GROMACS 2024.3 [23]. For all Martini CG simulations, two initial steepest descent minimizations of each 5000 steps were performed (first with soft potential). Afterward, five successive equilibration steps were carried out in the NPT ensemble employing the v-rescale thermostat [24] with a reference temperature of T= 310 K and the Berendsen barostat [25] with a reference pressure of p = 1 bar with timesteps of 2, 5, 10. 15, 20 fs in which the restraints on the positions of the lipid headgroups in the Z direction of initially 200 *kJmol*^−1^*nm*^2^ were gradually released. The equilibration was, in total, 36 ns long. The cutoff distance for nonbonded interactions (van der Waals and Coulomb interactions) was set to 1.1 nm, and Coulomb interactions were treated with a reaction field. The time step was set to 20 fs during the production runs. The LINCS algorithm was used to fix all bond lengths. During the production runs, the Berendsen barostat was replaced by a Parrinello–Rahman barostat [26].

#### 3.1.2 Analysis

MD trajectories were analyzed using GROMACS 2024.3 [23], MDAnalysis [27], LiPyphilic package [28], and in-house scripts.

##### flip-flop count

The number of flip-flops was counted using custom Python scripts and MDAnalysis. First, the distance between the Fe bead of Fe-MBT and the center of the membrane in the Z direction was calculated. Then, the z-distance was classified into three regions: near upper leaflet headgroup, buffer zone, and near lower leaflet headgroup. The buffer zone was defined as *±*1.0 nm from the center of the membrane. A change in the leaflet is only counted as a flip-flop if a Fe-MBT molecule crosses the buffer zone, enters the other leaflet, and stays there for at least 20 ns. The flipping rate was calculated by the total number of successful flip-flops divided by the total simulation time and the number of Fe-MBT molecules in the system.

##### order parameter

The coarse-grained order parameter, as defined in Seo et al. [29], was calculated by the order parameter module from LiPyphilic package [28]. The coarse-grained order parameter is defined as:

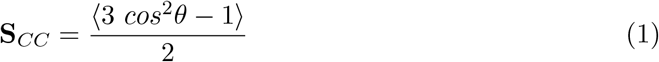

Where *θ* is the angle between the membrane normal and the vector connecting two consecutive tail beads, angular brackets denote averages over all beads in an acyl tail. For the lipid molecules, the order parameter was calculated as the weighted average of both acyl tails.

## 4 Results

### 4.1 Spantanous flipping of Fe-MBT is lipid specific

With the optimized ligand model, we investigated its behavior in various lipid membranes. Up to 100 Fe-MBT molecules (5 % of membrane composition) were placed in the differ- ent membranes, and two replicas for each were run. The composition and extent of the simulation runs are listed in Table 1.

Our simulations reveal a striking dependence of siderophore flip-flop dynamics on the lipid composition of the membrane. In bilayers composed of general bacterial lipids, such as palmitoyl-oleoyl-phosphatidylethanolamine (POPE), palmitoyl-oleoyl-phosphatidylglycerol (POPG), and cardiolipin (CL) the spontaneous translocation of the siderophore occurs rapidly, within the microsecond timescale. The flipping rates observed in these systems range from 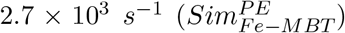 to 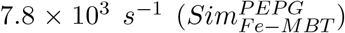, with multiple translocation events detected during the simulation time (Table 1). These results suggest that such lipid environments provide a highly permissive landscape for flip-flops, likely due to their intrinsic biophysical properties, such as membrane packing and hydration dynamics.

In contrast, mycobacterial-specific lipids, including mycobacterial phosphatidylethanolamine (TBPE), phosphatidylinositol (TBPI), and phosphatidylmyoinositol mannosides (PIMs), impose significant barriers to spontaneous flip-flop. In these membranes, translocation events are either not observed within the simulation timescale or occur as rare events on the millisecond scale—almost an order of magnitude slower than in membranes composed of generic bacterial lipids. For example, in the MtbPE and MtbMEMB systems, the flipping rate is reduced to 2.7 × 10^2^ *s*^−1^, and in the MtbPEPI system, no flip-flop was detected. Even in mixed mycobacterial/generic lipid compositions, such as MtbPE+POPG, the rate remains lower 1.2 × 10^3^ *s*^−1^), indicating that mycobacterial lipids significantly hinder translocation (Table 2).

**Table 2:**
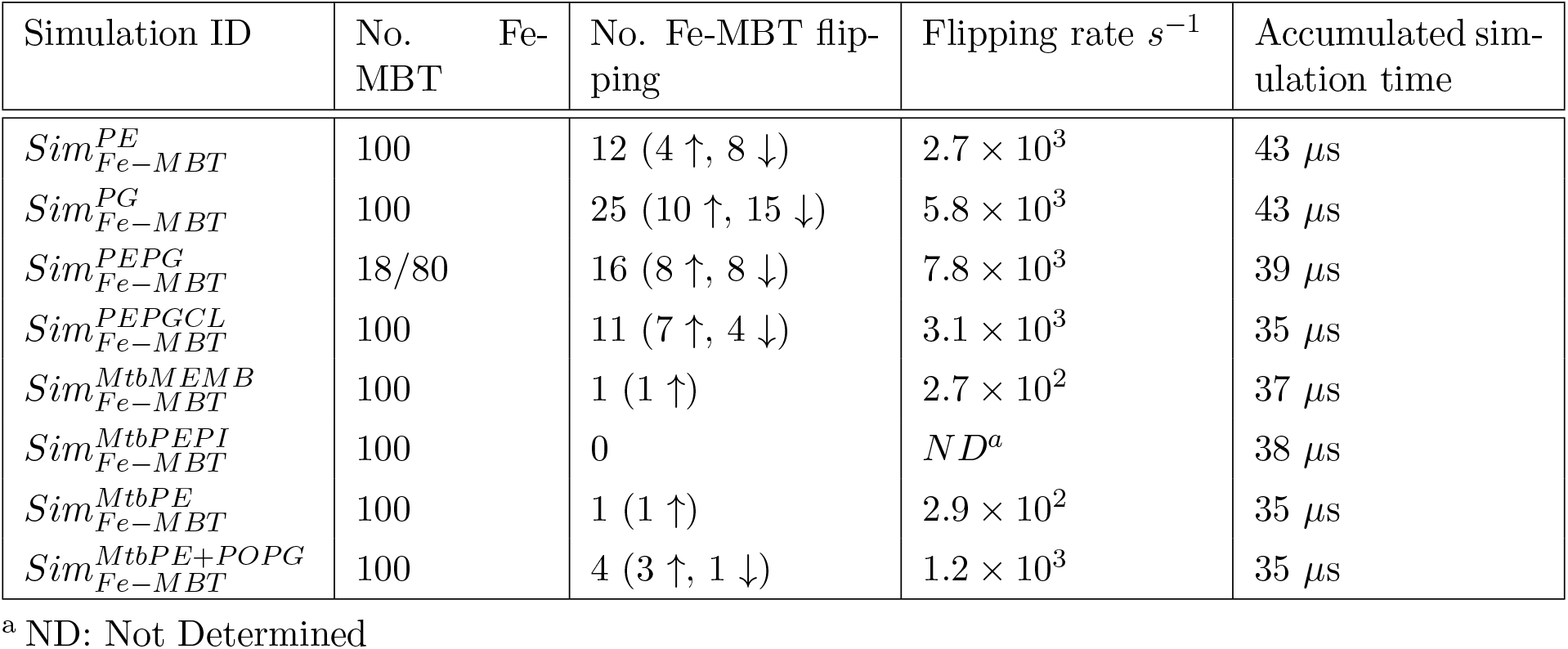
Analyzing the spontaneous flip-flop of the siderophore in membranes with different lipid compositions.

| Simulation ID | No. Fe-MBT | No. Fe-MBT flip-ping | Flipping rate $\text{s}^{-1}$ | Accumulated simulation time |
| --- | --- | --- | --- | --- |
| $Sim_{Fe-MBT}^{PE}$ | 100 | 12 (4 ↑, 8 ↓) | $2.7 \times 10^3$ | 43 $\mu\text{s}$ |
| $Sim_{Fe-MBT}^{PG}$ | 100 | 25 (10 ↑, 15 ↓) | $5.8 \times 10^3$ | 43 $\mu\text{s}$ |
| $Sim_{Fe-MBT}^{PEPG}$ | 18/80 | 16 (8 ↑, 8 ↓) | $7.8 \times 10^3$ | 39 $\mu\text{s}$ |
| $Sim_{Fe-MBT}^{PEPGCL}$ | 100 | 11 (7 ↑, 4 ↓) | $3.1 \times 10^3$ | 35 $\mu\text{s}$ |
| $Sim_{Fe-MBT}^{MtbMEMB}$ | 100 | 1 (1 ↑) | $2.7 \times 10^2$ | 37 $\mu\text{s}$ |
| $Sim_{Fe-MBT}^{MtbPEPI}$ | 100 | 0 | ND<sup>a</sup> | 38 $\mu\text{s}$ |
| $Sim_{Fe-MBT}^{MtbPE}$ | 100 | 1 (1 ↑) | $2.9 \times 10^2$ | 35 $\mu\text{s}$ |
| $Sim_{Fe-MBT}^{MtbPE+POPG}$ | 100 | 4 (3 ↑, 1 ↓) | $1.2 \times 10^3$ | 35 $\mu\text{s}$ |

<sup>a</sup> ND: Not Determined

**Table 3:** Averaged order parameter of lipid species in different systems.

| Simulation ID | $PE^a$ | PG | Fe-MBT |
| --- | --- | --- | --- |
| $Sim_{Fe-MBT}^{PE}$ | $0.359 \pm 0.20$ | – | $0.259 \pm 0.29$ |
| $Sim_{Fe-MBT}^{PG}$ | – | $0.337 \pm 0.21$ | $0.197 \pm 0.30$ |
| $Sim_{Fe-MBT}^{PEPG}$ | $0.354 \pm 0.20$ | $0.350 \pm 0.21$ | $0.237 \pm 0.29$ |
| $Sim_{Fe-MBT}^{PEPGCL}$ | $0.350 \pm 0.21$ | $0.354 \pm 0.21$ | $0.257 \pm 0.29$ |
| $Sim_{Fe-MBT}^{MtbMEMB}$ | $0.373 \pm 0.19$ | – | $0.324 \pm 0.27$ |
| $Sim_{Fe-MBT}^{MtbPEPI}$ | $0.415 \pm 0.19$ | – | $0.315 \pm 0.27$ |
| $Sim_{Fe-MBT}^{MtbPE}$ | $0.410 \pm 0.19$ | – | $0.317 \pm 0.27$ |
| $Sim_{Fe-MBT}^{MtbPE+POPG}$ | $0.365 \pm 0.19$ | $0.370 \pm 0.21$ | $0.261 \pm 0.29$ |

<sup>a</sup> POPE for general systems and TBPE for mycobacterial membranes.

These findings highlight the critical role of membrane composition in regulating siderophore dynamics. The restrictive nature of mycobacterial-specific lipids suggests a functional adaptation that could influence siderophore availability and transport, potentially affecting iron acquisition in mycobacteria.

The transition is relatively fast when a flip-flop occurs, typically within 100 ns (Figure 2). In some cases, translocation happens on even shorter timescales, in the order of just a few nanoseconds. This suggests that once the siderophore overcomes the initial energetic barrier to dissociate for one leaflet, its movement across the membrane to the other leaflet is rapid. The observed rapid transition further supports the idea that membrane composition primarily governs the rate-limiting step, likely by modulating the free energy barrier associated with dissociating the siderophore from the primary leaflet during translocation.

**Figure 2:**
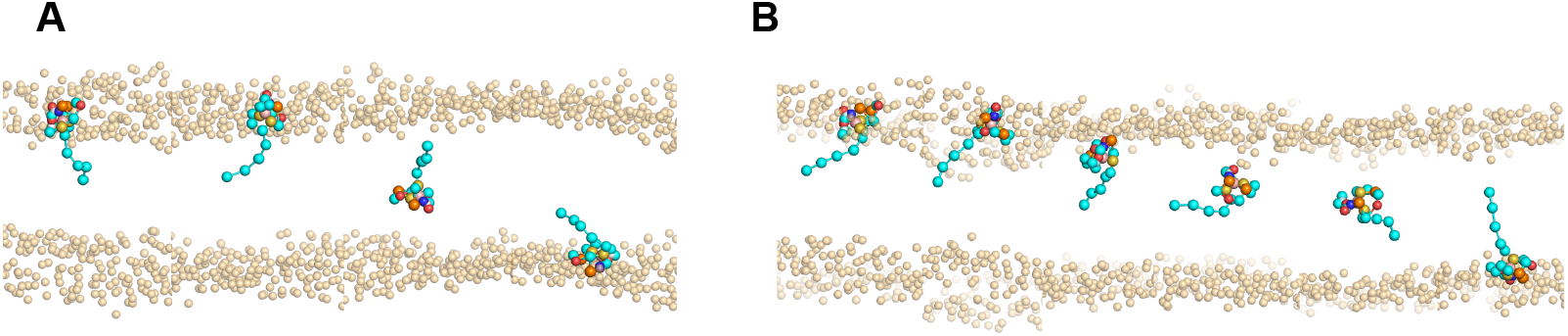
Spantanous flip-flop of Fe-MBT. Flipping in **A**) POPE environment 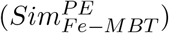. **B**) POPG environment 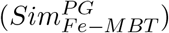.

To further understand the differences in flip-flop rates, we analyzed the order parameter of the lipid tails in each membrane system (Table 2). The results clearly distinguish between the general bacterial lipid membranes and the mycobacterial-specific lipid environments. In POPE- and POPG-containing bilayers, the order parameter of lipid acyl chains remains in the range of 0.337–0.359, indicating a relatively disordered state that likely facilitates the spontaneous translocation of the siderophore. In contrast, mycobacterial membranes, particularly those composed of MtbPE, MtbPEPI, and MtbMEMB, exhibit significantly higher order parameters (0.373–0.415), reflecting a more rigid and tightly packed membrane environment.

This increased lipid order in mycobacterial membranes correlates with the observed slower flip-flop rates (Table 1), suggesting that the tighter packing imposes a greater energetic barrier for the siderophore to translocate across the bilayer. Notably, the Fe-MBT order parameter follows the same trend, with values increasing from 0.197–0.259 in general lipid membranes to 0.315–0.324 in mycobacterial-specific environments. This shift indicates that the siderophore experiences a more restricted and less dynamic environment in mycobacterial membranes, further explaining the significant reduction in flip-flop frequency.

## 5 Discussion

The results of this study highlight the crucial role of membrane lipid composition in governing the spontaneous flip-flop of mycobacterial siderophores. Our simulations demonstrate that in membranes composed of general bacterial lipids such as POPE and POPG, translocation occurs readily within the microsecond timescale, facilitated by a relatively disordered lipid environment. This finding aligns with previous studies showing that lipid disorder and membrane fluidity strongly influence transbilayer diffusion of amphiphilic molecules [30]. In contrast, mycobacterial-specific lipids, including MtbPE, MtbPI, and PIMD, severely restrict siderophore flip-flop, with translocation events occurring nearly an order of magnitude slower or being entirely absent within our simulation timescales. These observations are consistent with prior work indicating that mycobacterial membranes exhibit unique biophysical properties, including low permeability and high rigidity [31, 32].

A key mechanistic insight from our analysis is the direct correlation between lipid tail ordering and flip-flop rates. The order parameter measurements reveal that mycobacterial membranes exhibit significantly higher lipid tail ordering than general bacterial membranes, suggesting a more rigid and tightly packed bilayer. This increased order likely imposes a higher energetic barrier for the siderophore to undergo spontaneous translocation. Similar trends have been reported in studies on sterol-containing membranes, where lipid packing density was shown to modulate the rate of small molecule diffusion across bilayers [33]. Furthermore, the siderophore exhibits higher order parameter in mycobacterial membranes, indicating that its dynamics are more restricted in these environments. These findings provide a mechanistic explanation for the observed differences in flip-flop rates and further support the idea that lipid composition is a key determinant of membrane transport properties.

The biological implications of these findings are particularly relevant to mycobacterial iron acquisition and siderophore transport. Mycobacteria rely on specialized siderophores such as mycobactins to scavenge iron from the host environment, and their ability to translocate across the membrane is crucial for effective iron uptake. The rigid nature of mycobacterial membranes may serve as a regulatory mechanism, slowing down uncontrolled siderophore movement and ensuring controlled uptake via dedicated transporter systems such as IrtAB [12]. Interestingly, a sliding mechanism has recently been proposed for flipping Fe-MBT by IrtAB [13]. It would be interesting to see whether the sliding on the surface of IrtAB is also affected by the rigidity of the mycobacterial membrane. This adaptation could be essential for maintaining iron homeostasis in the complex intracellular environment of pathogenic mycobacteria, where iron availability is tightly restricted by host defense mechanisms [34, 35].

While these results offer valuable insights, it is important to consider certain limitations of our study. First, our simulations are based on coarse-grained molecular dynamics, which efficiently sample long-timescale lipid dynamics but may oversimplify specific molecular interactions, such as hydrogen bonding or steric effects, that could influence siderophore translocation. A more detailed understanding of these mechanisms would require atomistic simulations, which, although computationally expensive, could provide a more accurate representation of the energetic barriers involved. Second, the composition of the mycobacterial membrane used in this study is based on the lipid species used by Brown et al. [36]. While this dataset captures the key lipid species, it does not fully reflect the diversity of mycobacterial lipidomes. Mycobacterial membranes are known to contain a mixture of saturated and unsaturated lipids, and the included lipids here primarily include more saturated lipid tails, which likely contribute to the high packing density observed in our simulations. However, other studies suggest that mycobacteria also incorporate a significant fraction of unsaturated lipids, which could lead to greater membrane fluidity than what is captured here. Future studies should include a more heterogeneous lipid composition, including unsaturated mycobacterial lipids, to assess how lipid tail flexibility influences siderophore translocation.

In conclusion, our findings provide molecular-level insights into how lipid composition influences siderophore translocation in bacterial membranes. Mycobacterial membranes’ distinct lipid ordering properties create a more restrictive environment, significantly reducing spontaneous flip-flop rates compared to general bacterial lipids. These results underscore the importance of membrane biophysical properties in modulating the dynamics of essential metabolites and suggest potential avenues for targeting mycobacterial iron uptake through membrane interactions. Future studies combining coarse-grained and atomistic simulations and experimental validation will be crucial to elucidate further the energetic barriers and structural determinants governing siderophore transport in mycobacteria.

## 6 Acknowledgments

We thank Sebastian Thallmair for the helpful advice regarding the development of ligand FF. The VSC (Flemish Supercomputer Center) and the EuroHPC supercomputer LUMI provided the high-performance computing resources and services used for performing the simulations in this work. A.R.M. acknowledges research support from Ghent University (BOF starting grant no. BOF.STG.2021.0037.01).

## Notes

### Competing Interest Statement

The authors have declared no competing interest.

