## Supplementary figures and tables for "Effect of membrane composition on interaction and permeation of a mycobacterial siderophore"

### 7 Supplementary materials

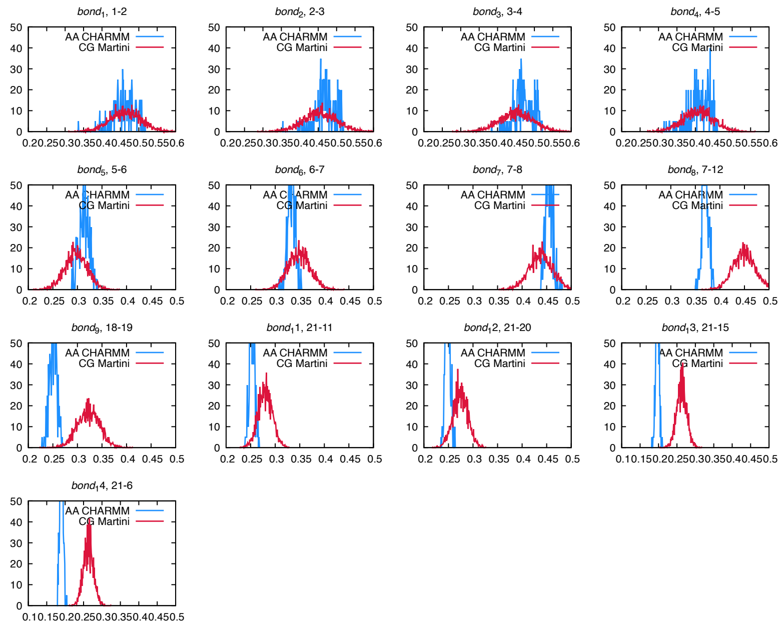


Figure S 1: **Bond distance distributions Fe-MBT CG model**. All-atom distributions are displayed in blue for the CHARMM-mapped trajectories, and Martini coarse-grained distributions are displayed in red

.

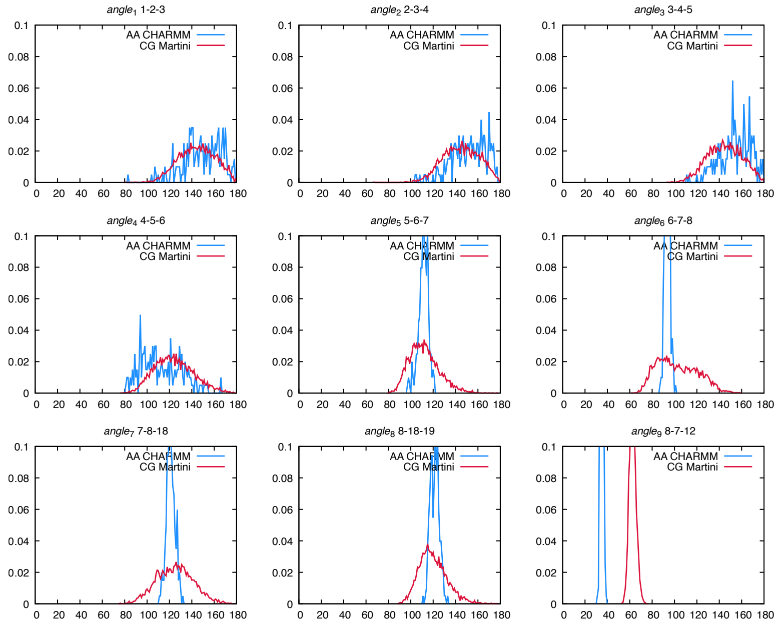


Figure S 2: **Angle distance distributions Fe-MBT CG model**. All-atom distributions are displayed in blue for the CHARMM-mapped trajectories, and Martini coarse-grained distributions are displayed in red

.

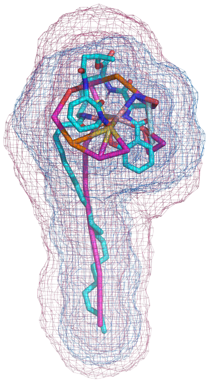

Figure S 3: **Solvent Accessible Surface Area (SASA) for Fe-MBT**. The blue wireframe represents the $CG_{SASA}$, and the red wireframe is the $AA_{SASA}$.
